# A quantitative trait variant in *Gabra2* underlies increased methamphetamine stimulant sensitivity

**DOI:** 10.1101/2021.06.29.450337

**Authors:** Lisa R. Goldberg, Emily J. Yao, Julia C. Kelliher, Eric R. Reed, Jiayi Wu Cox, Cory Parks, Stacey L. Kirkpatrick, Jacob A. Beierle, Melanie M. Chen, William E. Johnson, Gregg E. Homanics, Robert W. Williams, Camron D. Bryant, Megan K. Mulligan

**Affiliations:** Laboratory of Addiction Genetics, Department of Pharmacology and Experimental Therapeutics and Psychiatry, Boston, MA USA; NIGMS T32 Ph.D. Training Program in Biomolecular Pharmacology, Boston University School of Medicine, Boston, MA USA; Ph.D. Program in Bioinformatics, Boston University, Boston, MA USA; Program in Biomedical Sciences, Graduate Program in Genetics and Genomics, Boston University School of Medicine, Boston, MA USA; Department of Agricultural, Biology, and Health Sciences, Cameron University, Lawton, OK USA; Department of Medicine, Computational Medicine, Boston University School of Medicine; Departments of Anesthesiology, Neurobiology, and Pharmacology and Chemical Biology, University of Pittsburgh, Pittsburgh, PA USA; Department of Genetics, Genomics and Informatics, University of Tennessee Health Science Center, Memphis, TN USA

**Author notes:** co-senior authors, **Camron D. Bryant, Ph.D.**, Laboratory of Addiction Genetics, Department of Pharmacology and Experimental Therapeutics and Psychiatry, Boston University School of Medicine, 72 E. Concord St., L-606C, Boston, MA 02118 USA, E, **Megan K. Mulligan, Ph.D.**, Department of Genetics, Genomics and Informatics, 409 Translational Research Building, University of Tennessee Health Science Center, 71 South Manassas, Memphis, TN USA, E.

**Keywords:** cocaine, stimulant disorders, psychostimulant, addiction, amphetamine, methylphenidate, eQTL, quantitative trait gene, quantitative trait nucleotide

## Abstract

Psychostimulant (methamphetamine, cocaine) use disorders have a genetic component that remains mostly unknown. Here, we conducted genome-wide quantitative trait locus (QTL) analysis of methamphetamine stimulant sensitivity. To facilitate gene identification, we employed a Reduced Complexity Cross between closely related C57BL/6 mouse substrains and examined maximum speed and distance traveled over 30 min following methamphetamine (2 mg/kg, i.p.). For maximum methamphetamine-induced speed following the second and third administration, we identified a single genome-wide significant QTL on chromosome 11 that peaked near the *Cyfip2* locus [LOD = 3.5, 4.2; peak = 21 cM (36 Mb)]. For methamphetamine-induced distance traveled, we identified a single genome-wide significant QTL on chromosome 5 that peaked near a functional intronic indel in *Gabra2* that codes for the alpha-2 subunit of the GABA-A receptor [LOD = 5.2; peak = 35 cM (67 Mb)]. Striatal *cis*-expression QTL mapping corroborated *Gabra2* as a functional candidate gene underlying methamphetamine-induced distance traveled. CRISPR/Cas9-mediated correction of the mutant intronic deletion on the C57BL/6J background to the wild-type C57BL/6NJ allele was sufficient to reduce methamphetamine-induced locomotor activity toward the wild-type C57BL/6NJ-like level, thus validating the quantitative trait variant (QTV). These studies demonstrate the power and efficiency of Reduced Complexity Crosses in identifying causal genes and variants underlying complex traits. Functionally restoring *Gabra2* expression decreased methamphetamine stimulant sensitivity and supports preclinical and human genetic studies implicating the GABA-A receptor in psychostimulant addiction-relevant traits. Importantly, our findings have major implications for investigators studying psychostimulants in the C57BL/6J strain - the gold standard strain in biomedical research.

## INTRODUCTION

Psychostimulant (methamphetamine, cocaine) use disorders (**PUDs**) are a serious public health concern. Until the COVID-19 pandemic emerged, the opioid epidemic crisis had begun to plateau. Meanwhile, PUDs have quietly made a resurgence, with increased use and deaths (Cano *et al*. 2020; Maxwell 2020). Yet, despite an estimated 40-50% heritability for PUDs (Goldman *et al*. 2005; Ho *et al*. 2010; Ducci & Goldman 2012), genome-wide association studies have identified few loci (Jensen 2016). In one study, a significant GWAS hit for cocaine dependence mapped to ***FAM53B*** (Gelernter *et al*. 2014; Jensen 2016). Notably, an unbiased, quantitative trait locus (**QTL**) approach in mice identified a *trans*-expression QTL regulating *Fam53b* expression that was genetically correlated with variance in cocaine intravenous self-administration (**IVSA**) in BXD-RI mice, exemplifying cross-species bidirectional translation with discovery genetics in rodents.

**Reduced Complexity Crosses** exploit the extreme, near-isogenic nature of closely related inbred substrains to rapidly map, pinpoint, and validate quantitative trait loci (**QTLs**) containing causal quantitative trait genes (**QTGs**), and quantitative trait variants (**QTVs**) underlying complex trait variation (Bryant *et al*. 2018, 2020), including gene expression and behavior (Kumar *et al*. 2013; Kirkpatrick *et al*. 2017; Bryant *et al*. 2019; Mulligan *et al*. 2019). Of relevance to the present study, Kumar and colleagues used a mouse Reduced Complexity Cross between C57BL/6 substrains to map a missense variant in *Cyfip2* with sensitivity to cocaine-induced velocity and extended these findings to methamphetamine (Kumar *et al*. 2013). We previously used a similar Reduced Complexity Cross to map and validate *Cyfip2* in binge-like eating (Kirkpatrick *et al*. 2017). Also of relevance to the present study, we exploited the reduced complexity of C57BL/6 substrains to identify a functional noncoding single nucleotide deletion in *Gabra2* (alpha-2 subunit of the GABA-A receptor) that induced a loss-of-function decrease in transcript and protein expression (Mulligan *et al*. 2019). Correction of this mutation via CRISPR/Cas9 gene editing restored Gabra2 expression at both the transcript and protein levels (Mulligan *et al*. 2019). DBA/2 mouse substrains combined with historical BXD-RI substrains have also been exploited to identify a functional missense variant in *Taar1* (trace amine-associated receptor 1) underlying differences in the aversive properties of methamphetamine self-administration, body temperature (Harkness *et al*. 2015) and toxicity (Shi *et al*. 2016; Miner *et al*. 2017; Reed *et al*. 2017).

Administration of addictive drugs such as opioids and psychostimulants increases dopamine release in forebrain regions, including the dorsal striatum and nucleus accumbens, which contributes to the locomotor stimulant and rewarding properties of drugs of abuse (Adinoff, 2004; Di Chiara and Imperato, 1988). Psychostimulant-induced locomotor activity is a rapid, high-throughput heritable trait that is amenable to QTL mapping in multiple genetic populations (Phillips *et al*. 2008; Bryant *et al*. 2012a; Parker *et al*. 2016; Gonzales *et al*. 2018) and has a shared genetic basis with other addiction-relevant behavioral traits. As two examples, we mapped and validated genetic factors influencing psychostimulant and opioid-induced locomotor activity, including *Csnk1e* (Bryant *et al*. 2012c) and *Hnrnph1* (Yazdani *et al*. 2015). Subsequently, we and others have extended the role of these two genes to other complex behavioral models for addiction, including reward as measured via conditioned place preference (Goldberg *et al*. 2017; Ruan *et al*. 2020a) and reinforcement as measured via intravenous and oral self-administration (Wager *et al*. 2014; Ruan *et al*. 2020a).

C57BL/6J (B6J) and C57BL/6NJ (B6NJ) are two substrains of C57BL/6, the most commonly used mouse strain in biomedical research, and are 99.9% genetically similar, yet exhibit significant differences in several addiction-associated traits (Bryant *et al*. 2008), including ethanol consumption (Mulligan *et al*. 2008; Jimenez Chavez *et al*. 2021), nicotine behaviors (Akinola *et al*. 2019), and psychostimulant behaviors (Bryant *et al*. 2008; Kumar *et al*. 2013). Although phenotypic differences between B6 substrains can be quite large, genotypic diversity is extremely small, with only an estimated 10,000 to 20,000 variants (SNPs plus indels) distinguishing the two strains (Keane *et al*. 2011; Yalcin *et al*. 2011; Simon *et al*. 2013).

In the present study, we used a Reduced Complexity Cross between C57BL/6 substrains to map the genetic basis of sensitivity to the locomotor stimulant properties of methamphetamine, including maximum speed and distance traveled. Following the identification of two historical loci, including one locus for sensitized methamphetamine-induced maximum speed near the *Cyfip2* missense mutation that was previously identified for acute and sensitized cocaine velocity (Kumar *et al*. 2013) and a second locus near the functional intronic variant in *Gabra2* (Mulligan *et al*. 2019), we used a CRISPR/Cas9 gene-edited knockin mouse model with the corrected *Gabra2* mutation to validate this functional indel as necessary for enhanced acute stimulant sensitivity that is exhibited in the parental C57BL/6J substrain (Kumar *et al*. 2013).

## MATERIALS AND METHODS

### C57BL/6J (B6J), C57BL/6NJ (B6NJ), and a B6J x B6NJ-F2 Reduced Complexity Cross (Bryant Lab, BUSM)

All experiments involving mice were approved by the Boston University and University of Tennessee Health Science Center Institutional Animal Use and Care Committees and were conducted in accordance with the AAALAC Guide for the Use and Care of Laboratory Animals (National Research Council (US) Committee for the Update of the Guide for the Care and Use of Laboratory Animals 2011). Mice were housed in an AAALAC-accredited temperature- and climate-controlled facilities on a 12 h light/dark cycle (lights on at 0630 h). Mice were housed in same-sex groups of two to five mice per cage with standard laboratory chow and water available ad libitum except during testing. Mice were 50-100 days old on the first day of training. All behavioral testing was performed during the light phase of the 12 h light/dark cycle (0800 h to 1300 h).

C57BL/6J (**B6J**; n=31) and C57BL/NJ (**B6NJ**; n=32) mice were purchased from The Jackson Laboratory (Bar Harbor, ME USA) at 7 weeks of age and were habituated in the vivarium one week prior to experimental testing that occurred next door. All behavioral testing was performed during the light phase of the 12 h light/dark cycle (0800 h to 1300 h). For QTL mapping, a unidirectional cross was conducted whereby B6J females were crossed to B6NJ males to generate B6J x B6NJ-F_1_ mice and B6J x B6NJ F_1_ offspring were intercrossed to generate B6J x B6NJ F_2_ mice. All mice within a cage were assigned the same treatment.

All mice comprising the parental substrains and F2 offspring that were behaviorally tested had a prior, identical history of naloxone-induced conditioned place aversion as described in our original publication (Kirkpatrick & Bryant 2015). Briefly, following initial assessment of preference for the drug-paired side, 24 h later, mice received two alternating injections of naloxone hydrochloride (4 mg/kg, i.p.) and two alternating injections of saline (i.p.), separated by 48 h. Seventy-two hours and 96 hours after the second saline trial, mice were re-assessed for drug-free and state-dependent conditioned place aversion for the naloxone-paired side, respectively. Thus, all mice received a total of three injections of 4 mg/kg naloxone over 9 days. One week following recovery from the test for naloxone-induced conditioned place aversion, mice were tested for methamphetamine stimulant sensitivity as described below.

### *Gabra2* knockin mice

Gene-edited knockin mice were generated by inserting the corrected *Gabra2* intronic nucleotide on the mutant C57BL/6J background via CRISPR/Cas9 gene editing as previously described (Mulligan *et al*. 2019). Briefly, a sgRNA was designed that targeted *Gabra2* at the intron/exon junction near chromosome 5 at 71,014,638 Mb (mm10). A T7 promoter containing the sgRNA template was used to produce sgRNA and Cas9 mRNA that was then purified, ethanol precipitated, and re-suspended in DEPC-treated water. A 121 nucleotide single stranded DNA repair template oligo with the T insertion in the intron of *Gabra2* along with sgRNA, and *Cas9* mRNA were co-injected into the cytoplasm of B6J one-cell embryos. Offspring of injected embryos were screened for the insertion via PCR amplification of the knockin site. PCR products containing the amplicon were sequenced directly or subcloned into pCR2.1-TOPO (Invitrogen) and sequenced. The male founder (F0) was crossed to female B6J mice to generate F1 progeny. F1 mice were crossed to generate F2 mice. The colony is maintained through heterozygous breeding and all behavioral phenotyping was performed in generations F2 and higher. Potential off-targets were screened using CRISPOR (Haeussler *et al*. 2016; Concordet & Haeussler 2018) as previously described (Mulligan *et al*. 2019) and no off-target modifications were detected in the top 15 predicted off-target sites.

### Drugs

Methamphetamine hydrochloride (MA) (Sigma, St. Louis, MO USA) was dissolved in sterilized, physiological saline (0.9%) prior to injection (10 ml/kg, i.p.). The dose of MA (2 mg/kg) was chosen based on our prior success in mapping QTLs with this dose (Bryant *et al*. 2012c, 2012b; Yazdani *et al*. 2015; Ruan *et al*. 2020b) and based on a previous study that identified C57BL/6 substrain differences in MA-induced locomotor activity (Kumar *et al*. 2013).

### Methamphetamine-induced maximum speed and distance traveled in B6J, B6NJ, and B6J x B6NJ-F2 mice (Bryant Lab, BUSM)

The plexiglas apparatus consisted of an open field (40 cm length x 20 cm width x 45 cm tall; Lafayette Instruments, Lafayette, IN, USA) surrounded by a sound-attenuating chamber (MedAssociates, St. Albans, VT, USA). Behaviors were recorded using a security camera system (Swann Communications, Melbourne, Australia) and then video tracked (Anymaze, Stoelting, Wood Dale, IL, USA). We employed a five-day locomotor protocol (Ruan *et al*. 2020b), which is an extended version of the three-day protocol comprising the acute methamphetamine response (Yazdani et al., 2015). On Days (D)1 and 2, mice were injected with SAL (10 ml/kg, i.p.). On D3, D4, and D5 mice were injected with methamphetamine (2 mg/kg, 10 ml/kg, i.p.). Following i.p. injection, mice were immediately placed into the open field and video recorded over 30 min. Locomotor phenotypes, including total distance traveled and maximum speed while mobile, were calculated with AnyMaze.

Data were analyzed using repeated measures ANOVA with Strain and Sex as factors and Day as a categorical repeated measure. Time course analysis for a given day was analyzed in a similar manner but with Time (six, 5-min bins) as the repeated measure. Post-hoc analysis was conducted using the family-wide error variance with simple contrasts and Tukey’s honestly significant difference to correct for multiple comparisons.

### Methamphetamine-induced distance traveled in Gabra2 knockin (KI) mice (Mulligan Lab, UTHSC)

Validation for the role of the *Gabra2* functional intronic single nucleotide deletion (see below) in methamphetamine-induced distance traveled was conducted in the Mulligan Lab at UTHSC. Procedures were similar, though not identical. Here, a larger open field arena (40 cm x 40 cm x 40 cm) was employed and there were no sound attenuating chambers. The larger arena size (1.5-fold larger than the arena used in the Bryant Lab) likely accounts for the overall higher level of locomotor activity in this study compared to the parental strain and F2 studies. Similar to the Bryant Lab, behavior was also recorded with a security camera system and locomotor phenotypes were also calculated in AnyMaze.

### DNA collection and genotyping in F2 mice

DNA was extracted from spleens of F2 mice and prepared for genotyping using a standard salting out protocol. Ninety SNP markers spaced approximately 30 Mb (approximately 15 cM) apart were genotyped using a custom-designed 96 x 96 Fluidigm SNPtype array (South San Francisco, CA USA) (Kirkpatrick *et al*. 2017; Bryant *et al*. 2019). SNPs were called using the Fluidigm SNP Genotyping Analysis Software and SNPtype normalization with the default threshold.

### QTL analysis in B6J x B6NJ-F2 mice

QTL analysis was performed in F_2_ mice using the R package R/qtl as previously described (Broman *et al*. 2003; Bryant *et al*. 2009; Kirkpatrick *et al*. 2017). Quality checking of genotypes and QTL analysis were performed in R (https://www.r-project.org/) using R/bestNormalize (https://github.com/petersonR/bestNormalize) and R/qtl (Broman *et al*. 2003). Phenotypes were assessed for normality using the Shapiro-Wilk Test. Because some of the data residuals deviated significantly from normality, we used the orderNorm function to perform Ordered Quantile normalization (Peterson & Cavanaugh 2019) on all phenotypes. QTL analysis was performed using the “scanone” function and Haley-Knott (HK) regression. Permutation analysis (perm = 1000) was used to compute genome-wide suggestive (p < 0.63) and significance thresholds for log of the odds (LOD) scores (p < 0.05). Sex was included as an additive covariate in the QTL model. Peak marker positions were converted from sex-averaged cM to Mb using the JAX Mouse Map Converter (http://cgd.jax.org/mousemapconverter). Percent phenotypic variance explained by each QTL was calculated using the “fitqtl” function.

Power analysis of Day 5 maximum speed and Day 3 distance traveled using the data from 184 F2 mice was conducted using the R package QTLdesign (Sen *et al*. 2007). For each phenotype, we generated plots showing power versus % variance explained for an additively inherited QTL.

### RNA-seq for expression QTL (eQTL) analysis

Striatum was chosen for eQTL anlaysis for historical reasons and because this brain region is a major local site of drug action where methamphetamine binds to the dopamine transporter (and other monoamine transporters) and vesicular monoamine transporters to cause reverse transport of dopamine into the synapse, thus inducing stimulant, rewarding, and reinforcing effects (Chiu & Schenk 2012). Furthermore, Gabra2-containing GABA-A receptors are concentrated in the striatum (Morris *et al*. 2008) and are a dominant receptor type (Schwarzer *et al*. 2001). Thus, the striatum is a highly relevant brain tissue to ascertain the effects of *Gabra2* genetic variation on the transcriptome as they relate to methamphetamine-induced locomotor stimulation.

Striatum punches were collected as described (Yazdani *et al*. 2015) for RNA-seq from 23 F2 mice. Details on the prior experimental history of these mice are published (Bryant *et al*. 2019). Mice were previously trained and tested for place conditioning to oxycodone hydrochloride (1.25 mg/kg, i.p.; a total of 3 injections over 9 days). Six days later, these mice underwent an additional 4 daily injections of oxycodone (20 mg/kg, i.p.) before being tested for antinociception on the hot plate and then the following week, an additional 4 daily injections of the same dose of oxycodone prior to testing for affective withdrawal on the elevated plus maze 16 h later. Brains were harvested 24 h after behavioral assessment of oxycodone withdrawal (approximately 40 h after the final injection of oxycodone).

Brains were rapidly removed and sectioned with a brain matrix to obtain a 3 mm thick section where a 2 mm diameter punch of the striatum was collected. Left and right striatum punches were pooled and immediately placed in RNAlater (Life Technologies, Grand Island, NY, USA) and stored for 48 h at 4°C prior to storage in a −80°C freezer. Total RNA was extracted using the RNeasy kit (Qiagen, Valencia, CA, USA) as described (Yazdani *et al*. 2015). RNA was shipped to the University of Chicago Genomics Core Facility for cDNA library preparation using the Illumina TruSeq (oligo-dT; 100 bp paired-end reads). Libraries were prepared according to Illumina’s detailed instructions accompanying the TruSeq^®^ Stranded mRNA LT Kit (Part# RS-122-2101). The purified cDNA was captured on an Illumina flow cell for cluster generation and sample libraries were sequenced at 23 samples per lane over 5 lanes (technical replicates) according to the manufacturer’s protocols on the Illumina HiSeq 4000 machine, yielding an average of 69.4 million paired-end reads per sample. FASTQ files were quality checked via FASTQC and possessed Phred quality scores > 30 (i.e. less than 0.1% sequencing error). This dataset is publicly available on Gene Expression Omnibus (GEO #119719).

### *Cis*- and *trans*-expression QTL (eQTL) analysis

Details on eQTL mapping are published (Bryant *et al*. 2019). We aligned FastQ files to the reference genome (mm38) via TopHat (Trapnell *et al*. 2012) using the mm38 build and Ensembl Sequence and genome annotation. We used *featureCounts* to count and align reads. For *cis*-eQTL analysis, we used the same marker panel that we used in behavioral QTL analysis. We removed lowly expressed exons that did not possess at least 10 reads total across all 115 count files. Because of the low resolution of QTL mapping in an F2 cross, we liberally defined a gene with a *cis*-eQTL as any gene possessing a genome-wide significant association between expression and a polymorphic marker that was within 70 Mb of a SNP (the largest distance between any two SNPs from the 90-SNP panel). Analysis was conducted using *limma* with default TMM normalization and VOOM transformation (Law *et al*. 2014; Ritchie *et al*. 2015). A linear model was employed whereby sample replicates were treated as a random effects repeated measure. The duplicateCorrelation() function was used to estimate within-sample correlation, which was then included in the lmFit() function. An ANOVA test was conducted for gene expression that included Sex as a covariate and Genotype as a fixed effect. Genelevel tests were conducted using the likelihood Ratio test. A false discovery rate of 5% was employed as the cut-off for statistical significance (Benjamini & Hochberg 1995).

### Enrichment analysis

Enrichment analysis of genes whose transcripts correlated with Gabra2 transcript levels (r ≤ −0.5 or r ≥ 0.5; p ≤ 0.015) was conducted using the online tool Enrichr (Chen *et al*. 2013; Kuleshov *et al*. 2016) where we report GO terms for molecular, cellular, and biological function.

## RESULTS

### B6J mice show an increase in methamphetamine-induced maximum speed and locomotor activity compared to B6NJ mice

In examining B6 substrain differences in maximum speed, there were no substrain difference in response to saline injections (10 ml/kg, i.p.) on Days 1 and 2 (p’s > 0.05). However, in response to methamphetamine (2 mg/kg, i.p.) on Day 3, Day 4, and Day 5, B6J mice showed increased maximum speed relative to B6NJ mice, regardless of whether the data were collapsed across Sex or analyzed separately in females and males (**Fig.1A-C**).

**Figure 1.**
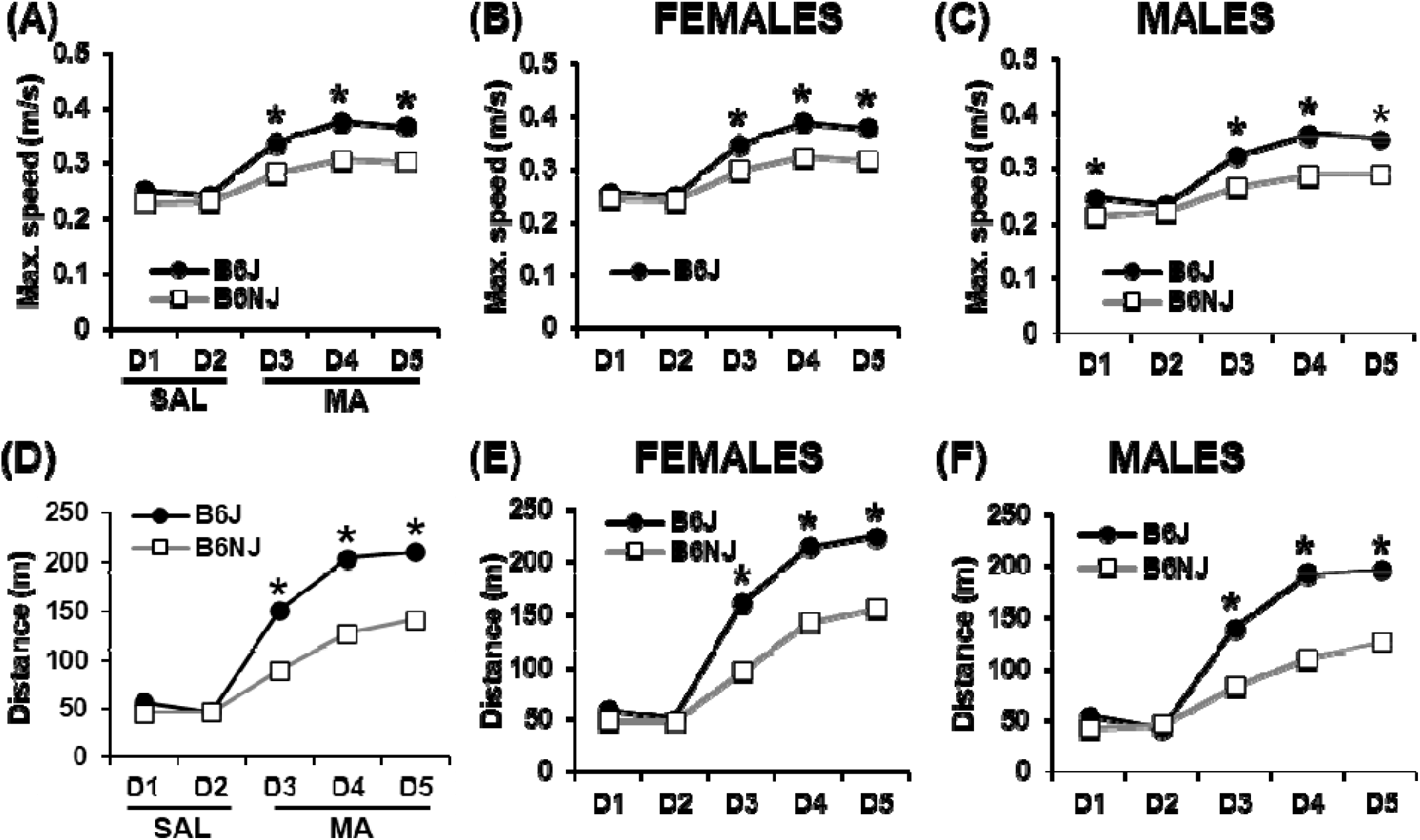
Maximum speed and distance traveled in response to saline (Days 1-2) and methamphetamine (Days 3-5) in the parental C57BL/6J (B6J) and C57BL/6NJ (B6NJ) substrains. **(A):** Sex-combined maximum speed (m/s) across training days. RM ANOVA across days indicated a main effect of Substrain (F_1,59_ = 59.96, p = 1.49 x 10^-10^), Sex (F_1,59_ = 18.02, p = 7.85 x 10^-5^), Day (F_4, 236_ = 377.90, p < 2 x 10^-16^), and a Substrain x Day interaction (F_4,236_ = 24.51, p < 2 x 10^-16^). Tukey’s post-hoc revealed a significant increase in maximum speed in B6J vs. B6NJ mice on Day(D) 3, D4, and D5 (*all p’s_adjusted_ < 0.0001). **(B):** Maximum speed (m/s) across training days in females. RM ANOVA indicated a main effect of Substrain (F_1,30_ = 22.29, p = 5.11 x 10^-5^), Day (F_4,120_ = 199.44, p < 2 x 10^-16^), and a Substrain x Day interaction (F_4,120_ = 13.21, p = 6 x 10^-9^). Tukey’s post-hoc test revealed a significant difference increase in maximum speed in B6J vs. B6NJ females on D3 (*p_adjusted_ = 0.0005), D4, and D5 (*p’s_adjusted_ < 0.0001). **(C):** Maximum speed across training days in males. RM ANOVA indicated a main effect of Substrain (F_1,29_ = 38.67, p = 8.75 x 10^-7^), Day (F_4,116_ = 178.66, p < 2 x 10^-16^), and a Substrain x Day interaction (F_4,116_ = 11.67, p = 5.36 x 10^-8^). Tukey’s post-hoc revealed a significant increase in maximum speed in B6J versus B6NJ males on D1 (p_adjusted_ = 0.03), D3, D4, and D5 (*p’s_adjusted_ < 0.0001). **(D):** Sex-combined distance traveled (m) across training days. RM ANOVA revealed a main effect of Substrain (F_1,59_ = 82.39, p = 8.51 x 10^-13^), Day (F_4,236_ = 798.0, p < 2 x 10^-16^), a Substrain x Day interaction (F_4,236_ = 67.99, p < 2 x 10^-16^), and a Sex x Day interaction (F_4,236_ = 6.46; p = 6.0 x 10^-5^). Tukey’s post-hoc revealed a significant increase in distance traveled in B6J vs. B6NJ mice on D3, D4, and D5 (all p’s_adjusted_ < 0.0001). **(E):** Distance traveled across training days in females. RM ANOVA indicated a main effect of Substrain (F_1,30_ = 41.18, p = 4.37 x 10^-7^), Day (F_4,120_ = 408,83, p < 2 x 10^-16^), and a Substrain x Day interaction (F_4,120_ = 27.05, p = 5.27 x 10^-10)^. Tukey’s post-hoc revealed a significant increase in distance traveled in B6J vs. B6NJ females on D3, D4, and D5 (*all p’s_adjusted_ < 0.0001). **(F):** Distance traveled across training days in males. RM ANOVA indicated a main effect of Substrain (F_1,29_ = 40.16, p = 6.34 x 10^-7^), Day (F_4,116_ = 396.98, p < 2 x 10^-16^), and a Substrain x Day interaction (F_4,116_ = 44.59, p < 2 x 10^-16^). Tukey’s post-hoc test revealed a significant increase in distance traveled in B6J vs. B6NJ males on Day 3, Day 4, and Day 5 (*all p’s_adjusted_ < 0.0001).

In examining B6 substrain differences in distance traveled following saline versus methamphetamine, there were no substrain differences in distance traveled in response to saline on Days 1 and 2 (p’s > 0.05). However, in response to methamphetamine, B6J mice showed an increase in distance traveled compared to B6NJ mice on Day 3, Day 4, and Day 5, regardless of whether the data were collapsed across Sex or analyzed separately in females and males (**Fig.1D-F**). These results replicate previous B6 substrain differences in the B6 mice from the J lineage show increased stimulant sensitivity compared to B6 mice from the N lineage (Kumar *et al*. 2013).

### Chromosome 11 QTL near the *Cyfip2* missense mutation underlying variation in sensitized methamphetamine-induced maximum speed (m/s)

Next, we sought to identify the genetic basis of differential methamphetamine-induced stimulant sensitivity in B6 substrains using an F2 Reduced Complexity Cross (Bryant *et al*. 2018, 2020). We identified a genome-wide significant QTL on chromosome 11 underlying maximum speed while mobile that was specific to methamphetamine treatment and emerged after the second and third methamphetamine injection on Day 4 and Day 5 (LOD = 3.5, 4.2; p = 0.039, 0.009; **Fig.2A**), thus requiring repeated methamphetamine exposure. The marker nearest the peak [rs48169870; 18 cM (31 Mb)] was located just proximally to the *Cyfip2* missense mutation [rs24064617401; 28 cM (46 Mb); **Fig.2B**] previously identified for acute and sensitized cocaine velocity (Kumar *et al*. 2013). Like the previous finding, the B6J allele was associated with increased methamphetamine-induced maximum speed (**Fig.2C**). Thus, the locus containing the *Cyfip2* missense mutation is associated with behavioral sensitivity to multiple psychostimulants.

**Figure 2.**
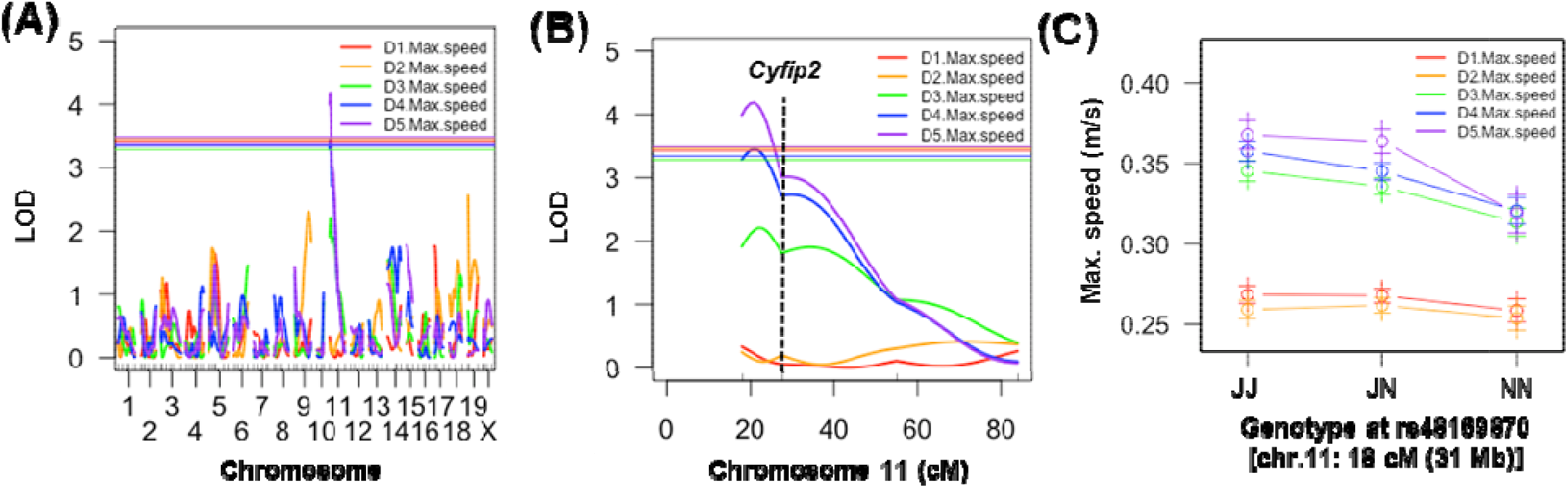
Genome-wide significant QTL on chromosome 11 near *Cyfip2* underlying variation in sensitized methamphetamine-induced maximum speed. Mice were treated on Day(D) 1 and D2 with saline (i.p.) and on D3, D4, and D5 with methamphetamine (2 mg/kg, i.p.) and behavioral activity was recorded over 30 min. **(A):** Genome-wide significant QTL on chromosome 11 for maximum speed following the second methamphetamine injection (2 mg/kg, i.p.) on D4 [LOD = 3.5, peak = 21 cM (36 Mb); Bayes interval: 18-39 cM (31-63 Mb); 11% of the phenotypic variance explained] and following the third methamphetamine injection (2 mg/kg, i.p.) on D5 [LOD = 4.2, peak = 21 cM (36 Mb); Bayes interval: 18-34 cM (31-57 Mb); 11% of the phenotypic variance explained]. Solid horizontal lines for panels A and B indicate significance thresholds for each phenotype (p < 0.05). **(B):** Chromosome 11 QTL plot for maximum speed on D1 through D5. **(C):** Effect plot of maximum speed as a function of Genotype at the peak locus for maximum speed on D1 through D5. J = homozygous for B6J allele; BN = heterozygous; N = homozygous for B6NJ allele.

### Chromosome 5 QTL near the *Gabra2* intronic deletion underlying variation in acute methamphetamine-induced locomotor activity (total distance traveled; m)

We identified two genome-wide significant QTLs on chromosome 5 that influenced locomotor activity (distance, m). The first QTL was for Day 2 distance traveled (saline) and was localized more proximally [peakmarker = 22 cM (41 Mb); LOD = 3.8; p = 0.037; **Fig.3A,B**]. The second QTL was located more distally [peak marker = 35 cM (67 Mb)] near the *Gabra2* intronic deletion (71 Mb) and was significant in response to the first and second methamphetamine injections on Day 3 and Day 4, respectively (LOD = 5.2, 3.6; p < 0.001, 0.034; **Fig.3A,B**). The B6J allele was associated with increased methamphetamine-induced distance traveled, as indicated by the effect plot in **Fig. 3C**. In examining the time course for the chromosome 5 QTL, the J allele was consistently associated with increased methamphetamine-induced distance traveled across the six, 5-min time bins (**Fig.3D**). Because we identified both a Sex x Day interaction and a Substrain x Day interaction in distance traveled in the parental substrains, we broke down and analyzed the time course of the effect plot separately in females and males and found that males clearly drove the QTL effect compared to females (**Fig.3E,F**).

**Figure 3.**
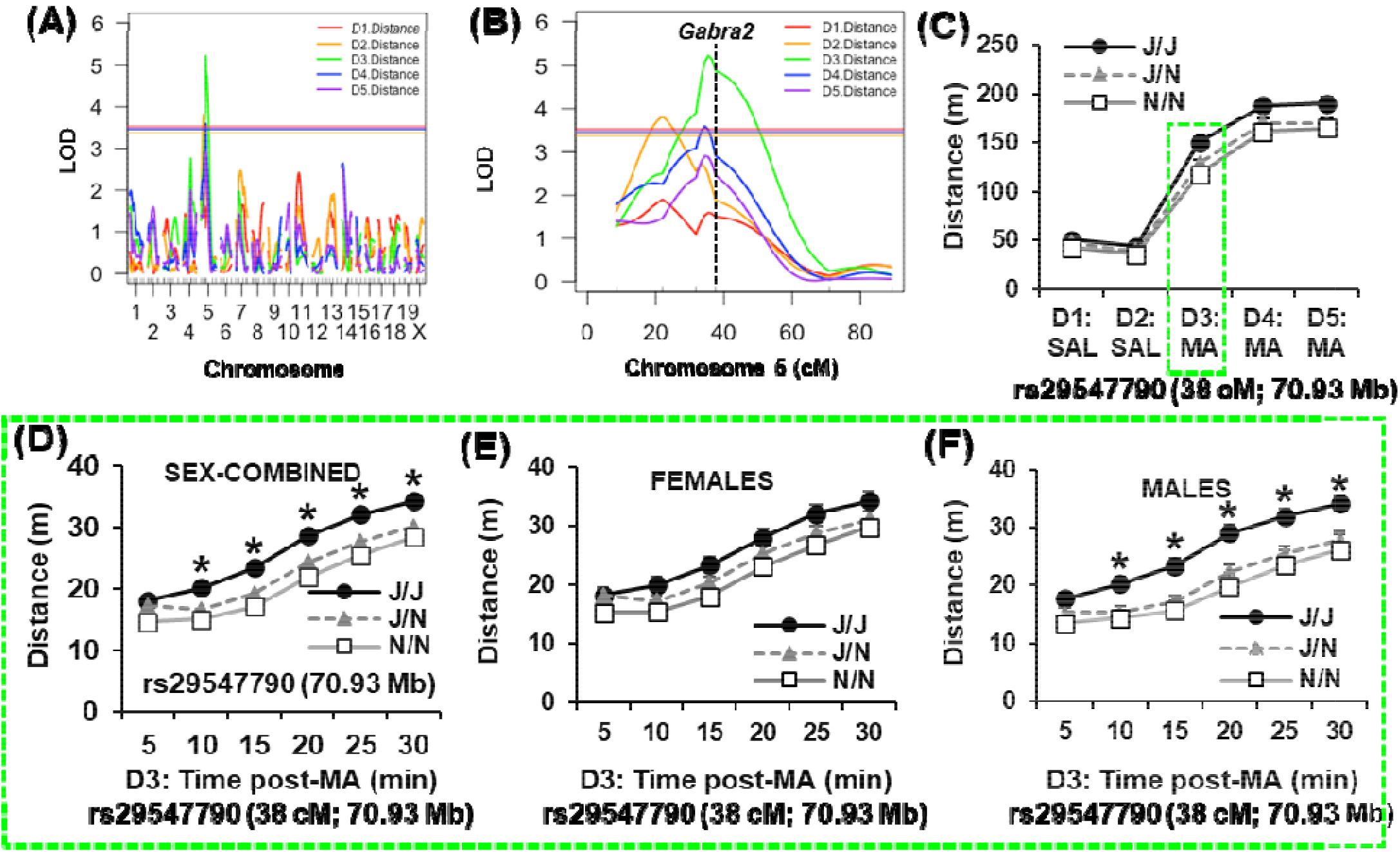
Genome-wide significant QTL on chromosome 5 near *Gabra2* underlying variation in acute methamphetamine-induced distance traveled. **(A):** Genome-wide significant QTL on chromosome 5 for distance traveled (m) on Day(D) 2 over 30 min following i.p. saline [LOD = 3.8, peak = 22 cM (41 Mb); Bayes interval: 22 cM (41 Mb); Bayes: 14-34 cM (29-66 Mb); 10% of the phenotypic variance explained], a second QTL for distance traveled D3 following 2 mg/kg methamphetamine [LOD = 5.2, peak = 35 cM (67 Mb); Bayes interval: 32-47 cM (60-95 Mb); 14% of the phenotypic variance explained], and a third QTL on D4 following the second methamphetamine injection [LOD = 3.6; peak = 34 cM (66 Mb); Bayes: 16-46 cM (30-93 Mb); 12% of the phenotypic variance explained]. Solid horizontal lines for panels A and B indicate significance threshold for each phenotype (p < 0.05). **(B):** Chromosome 5 QTL plot for distance traveled on D1 through D5. **(C):** Effect plot of total distance traveled for D1 through D5 over 30 min at the peak associated marker (rs29547790; 70.93 Mb). **(D):** Time course in 5-min bins of the effect plot for acute methamphetamine-induced distance traveled on D3. RM ANOVA indicated a main effect of Genotype (F_2,174_ = 7.20; p = 0.001), Sex (F_1,174_ = 3.89; p = 0.05), Time (F_5,870_ = 485.02; p < 2 x 10^-16^), a Genotype x Time interaction (F_10,870_ = 2.39; p = 0.0085) but no interactions with Sex (p’s > 0.46). * = significant increase in J/J relative J/N and N/N (Tukey’s p_adjusted_ < 0.05 for the three comparisons at each time point). **(E):** Time-course in females. RM ANOVA indiated no effect of Genotype (F_2,103_ = 2.13), an effect of Time (F_5,515_ = 290.22; p < 2 x 10^-16^), and no interaction (p = 0.51). Simple contrasts did not identify any significant differences among genotypes at any time point (p’s > 0.05). **(F):** Timecourse in males. There was a main effect of Genotype (F_2,71_ = 7.49; p = 0.0011), Time (F_5,355_ = 195.53; p < 2 x 10^-16^), and a Genotype x Time interaction (F_10,355_ = 1.85; p = 0.05). * = significant increase in J/J relative J/N and N/N (Tukey’s p_adjusted_ < 0.05 for the three comparisons at each time point). J = homozygous for B6J allele; BN = heterozygous; N = homozygous for B6NJ allele. Green, dashed traces denote distance traveled on D3.

Power analysis of the 184 F2 mice that we phenotyped in this study indicated that we had approximately 80% power to detect QTLs explaining at least 10% of the variance in maximum speed and in distance traveled (**Supplementary Figure 1**).

### Striatal *cis*-eQTL analysis identifies Gabra2 as the top transcript associated with rs29547790; the peak chromosome 5 marker linked to methamphetamine-induced distance traveled

Given that the likely causal variant underlying maximum speed induced by methamphetamine for the chromosome 11 locus was a missense variant in *Cyfip2* (Kumar *et al*. 2013), we turned our attention to the novel chromosome 5 locus whose peak marker (rs29547790; 70,931,531 bp) is located just proximally to a functional intronic deletion in *Gabra2* (71 Mb) (Mulligan *et al*. 2019) which codes for the alpha-2 subunit of the GABA-A receptor. In a genome-wide *cis*-eQTL analysis, we examined eQTLs associated with the peak chromosome 5 marker associated with methamphetamine-induced locomotor activity (rs29547790; 70.93 Mb). The top transcript was *Gabra2* (chromosome 5, 70.96 Mb; p_adjusted_ = 7.2 x 10^-27^; **Table 1**). Given this finding and the prior literature implicating Gabra2 in psychostimulant behavioral responses (Stephens *et al*. 2017), *Gabra2* is a top candidate quantitative trait gene underlying the chromosome 5 QTL for methamphetamine-induced locomotor activity. A list of 59 trans-eQTLs (p < 0.0001) showing a peak association with rs29547790 is provided in **Supplementary Table 2**.

**Table 1.** Cis-eQTL gene transcripts showing peak marker association in transcript variance with rs29547790 (chromosome 5: 70.9 Mb (FDR < 0.05)

| Gene | Chr.5 location (Mb) | Distance from rs29547790 (Mb) | Log2FC (JJ vs. NN) | P-value | Adjusted P-value |
| --- | --- | --- | --- | --- | --- |
| Gabra2 | 70.96 | 0.026 | -0.88 | 1.66E-30 | 7.24E-27 |
| Muc3a | 137.21 | 66.28 | 0.56 | 2.60E-10 | 1.14E-07 |
| Atp8a1 | 67.62 | 3.08 | 0.18 | 1.28E-06 | 9.57E-05 |
| Ptpn11 | 121.13 | 50.20 | 0.16 | 2.17E-05 | 0.000778 |
| Cxcl5 | 90.76 | 19.83 | -2.41 | 3.22E-05 | 0.001042 |
| Fgfr3 | 33.72 | 37.19 | 0.19 | 0.000123 | 0.002843 |
| Cux2 | 121.86 | 50.92 | -0.33 | 0.000136 | 0.003049 |
| Ankrd13a | 114.77 | 43.84 | 0.27 | 0.00029 | 0.005321 |
| Whsc1 | 33.82 | 37.03 | -0.14 | 0.000399 | 0.006869 |
| Mn1 | 111.42 | 40.49 | 0.24 | 0.000644 | 0.009812 |
| Auts2 | 131.44 | 60.51 | 0.20 | 0.000721 | 0.01069 |
| Cds1 | 101.77 | 30.83 | -0.09 | 0.002214 | 0.02399 |
| Lrrc8c | 105.52 | 34.59 | 0.22 | 0.002975 | 0.0293 |
| Tsc22d4 | 137.75 | 66.81 | 0.04 | 0.005113 | 0.043141 |
| Garem2 | 30.11 | 40.81 | 0.23 | 0.005705 | 0.046604 |

Red rows = decreased expression with the C57BL/6J allele versus the C57BL/6NJ allele; blue rows = increased expression with the C57BL/6J allele versus the C57BL/6NJ allele

**Table 2.** Enrichment analysis of genes correlated with Gabra2 transcript levels in F2 mice (n=23; r ≤ - 0.5 or r ≥ +0.5; p ≥ 0.015)

| **GO: BIOLOGICAL PROCESS** |  |  |  |  |  |  |  |
| --- | --- | --- | --- | --- | --- | --- | --- |
| Term | GO # | Overlap | P-value | Adj. P | Z-score | Combined Score | Genes |
| synaptic transmission, GABAergic | 51932 | 3/12 | 7.83E-05 | 0.05354 | -2.45 | 23.13 | GABRA2;GABRE;GABRG3 |
| GABA signaling pathway | 7214 | 3/20 | 3.89E-04 | 0.132938 | -2.06 | 16.21 | GABRA2;GABRE;GABRG3 |
| L-amino acid transport | 15807 | 3/26 | 8.59E-04 | 0.195793 | -2.22 | 15.68 | PRAF2;SLC3A2;SLC38A5 |
| regulation of postsynaptic membrane potential | 60078 | 3/34 | 1.89E-03 | 0.257775 | -1.51 | 9.49 | GABRA2;GABRE;GABRG3 |
| chemical synaptic transmission, postsynaptic | 99565 | 3/34 | 1.89E-03 | 0.257775 | -1.45 | 9.09 | GABRA2;GABRE;GABRG3 |
| **GO: CELLULAR COMPONENT** |  |  |  |  |  |  |  |
| Term | GO # | Overlap | P-value | Adj. P | Z-score | Combined Score | Genes |
| GABA-A receptor complex | 1902711 | 3/20 | 3.89E-04 | 0.023457 | -2.19 | 17.20 | GABRA2;GABRE;GABRG3 |
| dendrite membrane | 32590 | 3/21 | 4.51E-04 | 0.023457 | -2.78 | 21.38 | GABRA2;GABRE;GABRG3 |
| germ plasm | 60293 | 2/14 | 4.49E-03 | 0.123421 | -2.36 | 12.75 | TDRD1;SNRPG |
| cytoskeleton | 5856 | 10/521 | 4.75E-03 | 0.123421 | -1.58 | 8.45 | BCAS3;ACTR1A;EDA;MYOT;MVP;SHROOM2;MYOZ3;NPM3;S100A9;MDN1 |
| P granule | 43186 | 2/17 | 6.61E-03 | 0.137495 | -2.35 | 11.81 | TDRD1;SNRPG |
| **GO: MOLECULAR FUNCTION** |  |  |  |  |  |  |  |
| Term | GO # | Overlap | P-value | Adj. P | Z-score | Combined Score | Genes |
| benzodiazepine receptor activity | 8503 | 3/11 | 5.90E-05 | 0.009565 | -2.81 | 27.34 | GABRA2;GABRE;GABRG3 |
| GABA-gated chloride ion channel activity | 22851 | 3/13 | 1.01E-04 | 0.009565 | -2.49 | 22.93 | GABRA2;GABRE;GABRG3 |
| inhibitory<br>extracellular<br>ligand-gated<br>ion channel<br>activity | 5237 | 3/16 | 1.95E-04 | 0.012288 | -2.39 | 20.38 | GABRA2;GABRE;GABRG3 |
| ligand-gated<br>anion<br>channel<br>activity | 99095 | 3/19 | 3.32E-04 | 0.014028 | -2.16 | 17.29 | GABRA2;GABRE;GABRG3 |
| GABA-A<br>receptor<br>activity | 4890 | 3/20 | 3.89E-04 | 0.014028 | -2.09 | 16.42 | GABRA2;GABRE;GABRG3 |

### *Gabra2*-correlated transcripts and enrichment analysis

To gain insight into the neurobiological adaptations associated with differential *Gabra2* expression at the genomic level, we examine the correlation of Gabra2 with other transcripts genome-wide. We identified 148 transcripts with an absolute Pearson’s r value of 0.5 or greater (p ≤ 0.015; **Supplementary Table 3**). We conducted enrichment analysis of this gene list using Enrichr (Chen *et al*. 2013; Kuleshov *et al*. 2016) to identify GO molecular, cellular, and biological functions linked to decreased *Gabra2* expression. We identified several enrichment terms related to the GABA-A receptor signaling (**Table 2**) that were mainly driven by other GABA-A receptor subunits that were positively correlated with *Gabra2* expression (*Gabre, Gabrg3*; r = +0.56, +0.50; p ≤ 0.015; **Supplementary Table 4**). Other nominally, positively correlated GABA-A subunit transcripts included Gabrb1 (r = +0.37; p = 0.08) and Gabrq (r = +0.36; p = 0.09) (**Supplementary Table 4**). Additional GABA signaling-relevant transcripts that were correlated include the GPCR kinase coded by *Grk5* (r = +0.51; p = 0.013) (Kanaide *et al*. 2007) and the amino acid transporters coded by *Slc3a2* (r = −0.51; p = 0.014), and *Slc38a5* (r = −0.59; p = 0.003) (Cubelos *et al*. 2005).

**Table 3.**
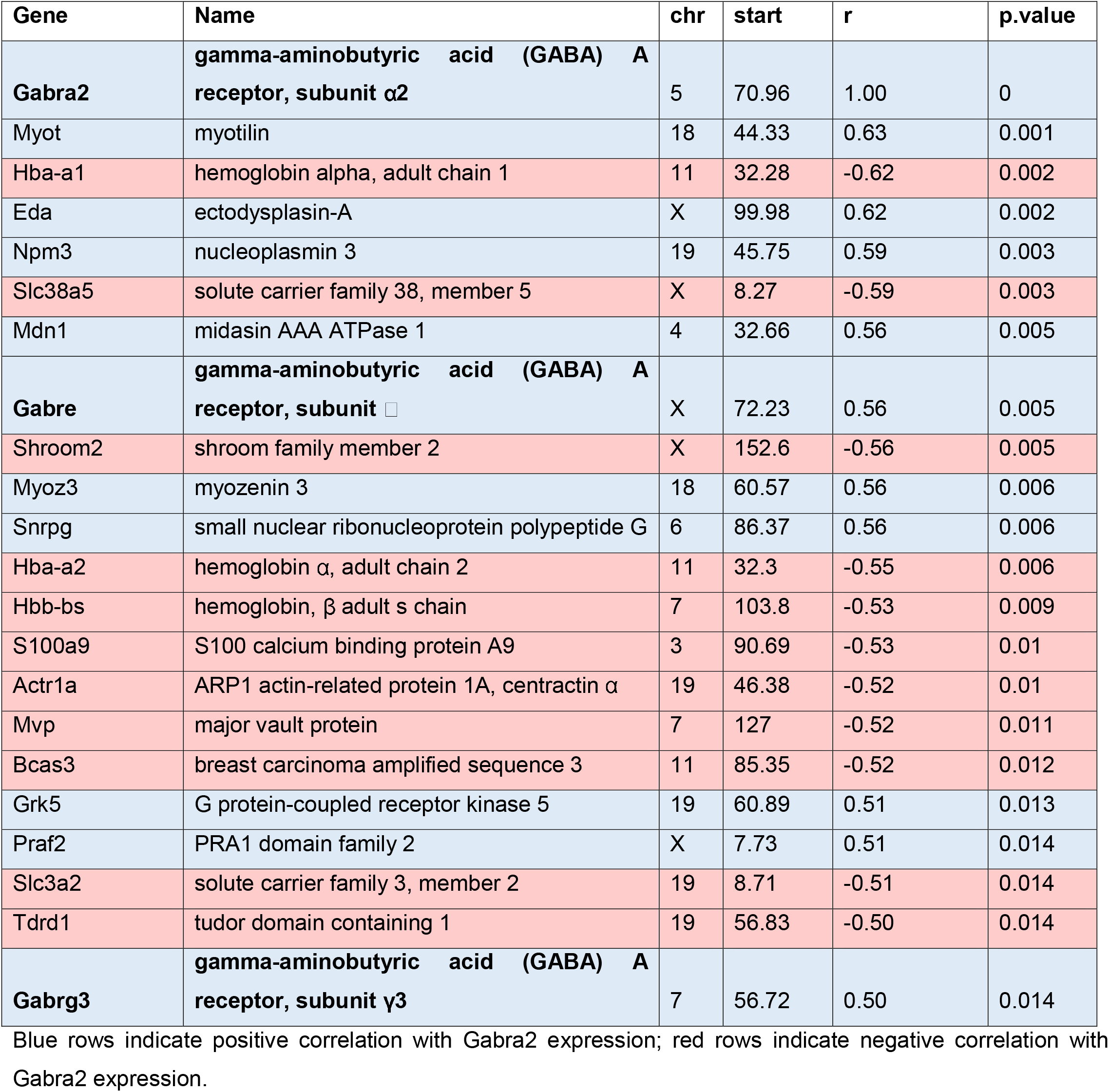
Pearson’s r for genes identified from enrichment analysis (see Table 2) that were correlated with Gabra2

### CRISPR/Cas9 correction of the functional intronic deletion in *Gabra2* reduces methamphetamine-induced distance traveled in B6J mice toward a B6NJ-like level

Based on co-mapping of a behavioral QTL with a highly robust striatal cis-eQTL for *Gabra2* expression and based on the known functional intronic variant within Gabra2 harbored by the B6J substrain that decreases Gabra2 expression (Mulligan *et al*. 2019), we tested the hypothesis that this single intronic nucleotide deletion in *Gabra2* located near a splice acceptor site (71,014,638 bp) in the B6J strain is the quantitative trait variant underlying variance in methamphetamine-induced locomotor activity. Specifically, we predicted that the *Gabra2* deletion increases methamphetamine sensitivity in B6J mice and that correction of this deletion via CRISPR-Cas9-mediated insertion of the B6NJ wild-type nucleotide would reduce methamphetamine-induced distance traveled toward a B6NJ-like level. Furthermore, given the more pronounced phenotypic effect of Genotype at the *Gabra2* locus in males compared to females (**Fig.3E, F**), we predicted that genotypic restoration of the wild-type allele would exert a more pronounced decrease in methamphetamine-induced distance traveled in male versus female mice.

Consistent with our predictions, there was no genotypic difference in distance traveled on Day 1 or Day 2 following saline (i.p.) injections. Sex-combined data from C57BL/6J mice homozygous for the Gabra2 mutational correction [reinsertion of the deleted intronic nucleotide; “knockin” (***Gabra2*** **KI**); **Fig.4A**] showed a decrease in methamphetamine-induced distance traveled on all three days of methamphetamine exposure (D3, D4, D5; **Fig.4B**). Consistent with the QTL results from F2 mice at the chromosome 5 locus, there was no genotypic difference in methamphetamine-induced locomotor activity in females on Day 3 (**Fig.4C**), while on Day 4 and Day 5, female KI mice showed a significant decrease in methamphetamine-induced locomotor activity (**Fig.4C**). On the other hand and again, consistent with the QTL results from F2 mice, male KI mice showed a significant decrease in methamphetamine-induced distance traveled following the first and second methamphetamine administration on Day 3 and Day 4, with no genotypic difference on Day 5 (**Fig.4D**).

**Figure 4.**
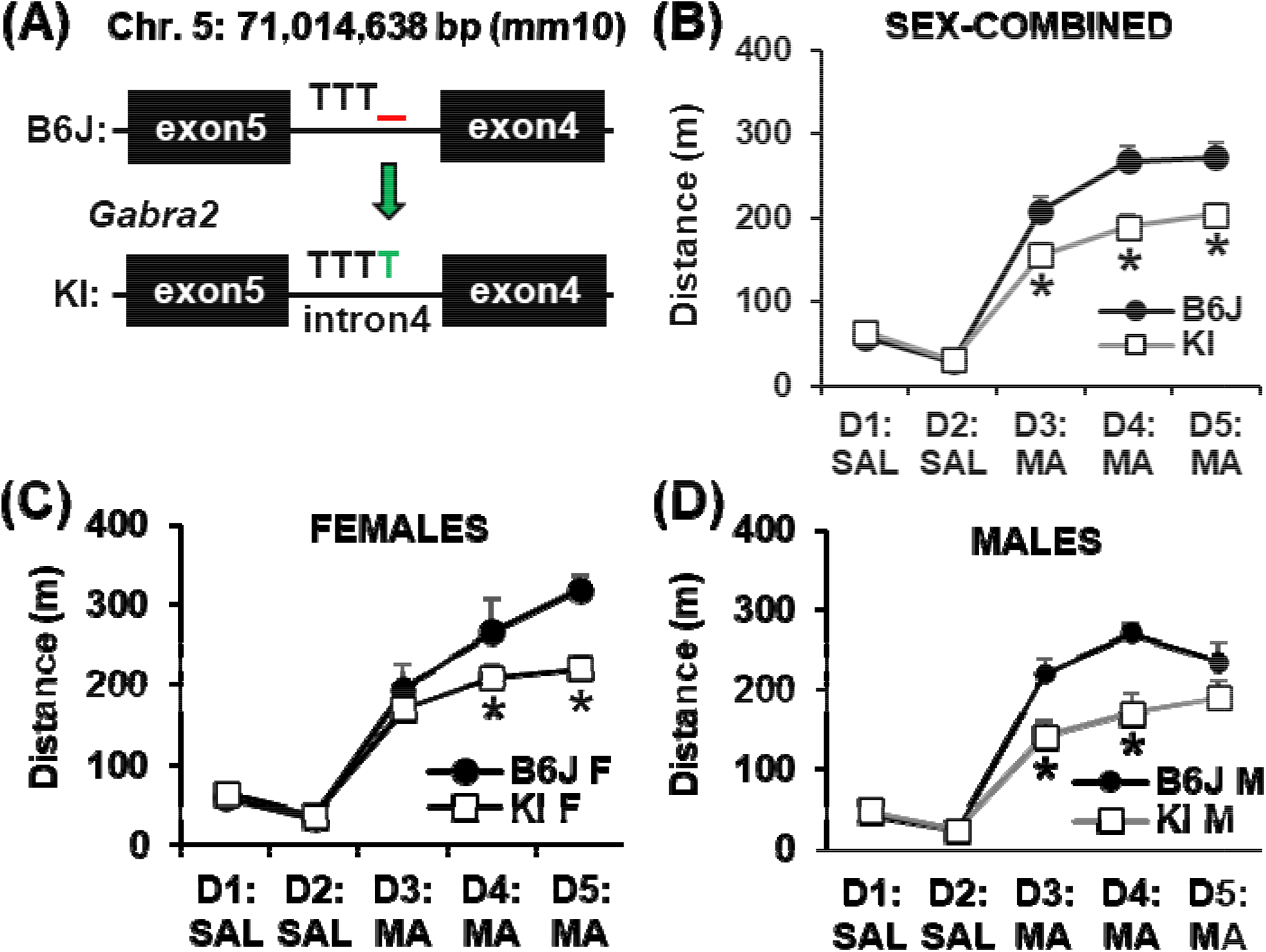
Identification of a quantitative trait variant in *Gabra2* that underlies variation in methamphetamine stimulant sensitivity as measured via distance traveled. **(A):** Schematic of gene-edited knockin (**KI**) of the single T nucleotide “corrected” allele that was inserted into intron 4 of the *Gabra2* gene. The C57BL/6J (B6J) substrain harbors a single nucleotide T deletion on chromosome 5 at 71,041,638 bp (mm10). CRISPR-Cas9 was used to insert the deleted T nucleotide onto the B6J genome thus “correcting” the single nucleotide deletion. **(B):** Distance traveled across Day(D) 1 through D5 in *Gabra2* KI vs. B6J wildtypes. RM ANOVA of the sex-combined data revealed a main effect of Genotype (F_1,31_ = 9.63, p = 0.0041), Day (F_4,124_ = 177.28, p < 2 x 10^-16^), and a Genotype x Day interaction (F_4,124_ = 7.36, p = 2.37 x 10^-5^). There was no statistically significant effect of Sex (F_1,31_ = 2.77; p = 0.11). Furthermore, the interaction of Sex with Day (F_4,124_ = 2.25; p = 0.068) and the Genotype x Sex x Day interaction (F_4,124_ = 2.09; p = 0.087) did not reach statistical significance. Simple contrasts of the sex-combined data revealed a significant decrease in distance traveled in KI vs. B6J mice on D3 (*p = 0.0063), D4 (*p < 0.0001), and D5 (*p = 0.0001). **(C):** Distance traveled in females. RM ANOVA indicated a main effect of Day (F_4,56_ = 73.86, p < 2 x 10^-16^) and a Genotype x Day interaction (F_4,56_ = 3.38, p = 0.015). Simple contrasts identified a significant decrease in distance traveled in *Gabra2* KI females vs. B6J wild-type females on D4 (*p = 0.0498) and D5 (*p = 0.0009). **(D):** Distance traveled in males. RM ANOVA indicated a main effect of Genotype (F_1,17_ = 6.49, p = 0.021), Day (F_4,68_ = 109.21, p < 2 x 10^-16^), and a Genotype x Day interaction (F_4,68_ = 6.60, p = 0.00015). Simple contrasts revealed a significant decrease in distance traveled in KI males vs. B6J wild-type males on D3 (*p = 0.0016) and D4 (*p = 0.0001), but not on D5 (p = 0.061).

Based on the QTL that was significant for the first methamphetamine exposure on Day 3 (**Fig.3A,B**; green traces), we were primarily interested in this phenotype and so we broke down the data for Day 3 into 5min bins like we did with the F2 mice (**Fig.3D-F**) to more closely examine the sex-combined and sex-stratified datasets. Similar to the F2 results for the chromosome 5 locus, the effect of Genotype on acute methamphetamine-induced locomotor activity was more pronounced in males. For the sex-combined dataset, *Gabra2* KI mice showed a significant decrease in methamphetamine-induced locomotor activity from 10 min to 30 min post-methamphetamine (**Fig.5A**). For females-only, there was no significant genotypic difference at any of the six time bins (**Fig.5B**). For males, there was a significant decrease in methamphetamine-induced locomotor activity in KI males from 15 min to 30 min (**Fig.5C**). Thus, like F2 males with the chromosome 5 QTL, male *Gabra2* KI mice account for the sex-combined phenotype of decreased methamphetamine-induced distance traveled in mice with the wild-type (KI) allele compared to the *Gabra2* deletion.

**Figure 5.**
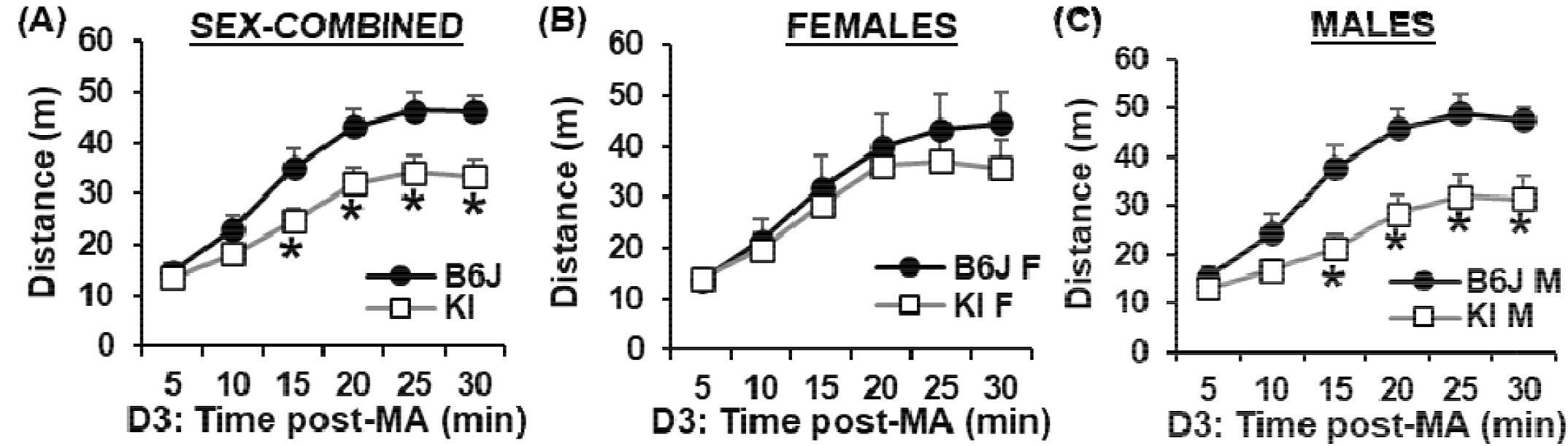
Time course of distance traveled following acute methamphetamine administration on Day 3 in *Gabra2* KI mice versus B6J wild-types. **(A):** Distance traveled on Day(D)3 across 5-min bins in KI vs. B6J wild-types. RM ANOVA revealed a main effect of Genotype (F_1,31_ = 5.723, p = 0.023), Time (F_5,155_ = 96.60, p < 2 x 10^-16^), and a Genotype x Time interaction (F_5,155_ = 4.27, p = 0.0011). Simple contrasts revealed a significant decrease in distance traveled at 15 min (p = 0.022), 20 min (p = 0.016), 25 min (p = 0.0078), and 30 min (p = 0.0046). **(B):** Time course of distance traveled in females on D3 following acute methamphetamine administration. RM ANOVA indicated a main effect of Time (F_5,70_ = 31.74, p < 2 x 10^-16^) but no effect of Genotype (F_1,14_ < 1) and no Genotype x Time interaction (F_5,70_ < 1). **(C):** Time course of distance traveled in males on D3 following acute methamphetamine administration. RM ANOVA indicated a main effect of Genotype (F_1,17_ = 8,32, p = 0.01), Time (F_5,85_ = 78.78, p < 2 x 10^-16^), and a Genotype x Time interaction (F_5,85_ = 6.53, p = 3.5 x 10^-5^). Simple contrasts revealed a significant decrease in KI males vs. B6J wild-type males at 15 min (*p = 0.0027), 20 min (*p = 0.0017), 25 min (*p = 0.0021), and 30 min (*p = 0.0034).

## DISCUSSION

We replicated the enhanced psychostimulant sensitivity in B6J compared to N substrains (Kumar *et al*. 2013) (**Fig.1**). We then used a Reduced Complexity Cross between C57BL/6 substrains to identify a QTL near the *Cyfip2* missense mutation previously identified for cocaine velocity (Kumar *et al*. 2013) that influenced methamphetamine-induced maximum speed (**Fig.2**). Next, we identified a novel QTL near the known functional intronic deletion in *Gabra2* (Mulligan *et al*. 2019) that influenced methamphetamine-induced distance traveled (**Fig.3**). *Cis*-eQTL analysis of striatal tissue from F2 mice implicated *Gabra2* as a causal quantitative trait gene underlying increased methamphetamine stimulant sensitivity (**Table 1**). Finally, CRISPR/Cas9 correction of the single nucleotide deletion via CRISPR-Cas9 reversed the enhanced methamphetamine sensitivity toward a C57BL/6NJ-like level (**Fig.4**), thus recapitulating the QTL effect (**Fig.5 vs. Fig.3**) and identifying the quantitative trait nucleotide.

Despite the fact that Gabra2 KI mice recapitulated the effect of Sex-dependent effect of Genotype at the chromosome 5 QTL containing the *Gabra2* deletion, neither result recapitulated the parental strain phenotype where both female B6J and male B6J mice harboring the *Gabra2* deletion showed qualitatively similar increases in methamphetamine-induced distance traveled on Day 3 (**Fig.1E,F**). This discrepancy is likely explained by the fact that these B6 substrains each harbor their own unique set of variants that contribute to the overall phenotype, including notably, the *Cyfip2* missense mutation in C57BL/6N that decreases psychostimulant velocity (Kumar *et al*. 2013).

To our knowledge, this is the first time that a *Gabra2* variant has been identified to influence methamphetamine behavior as previous studies focused primarily on cocaine, e.g., *Gabra2* knockouts with deletion of exon 4 (Dixon *et al*. 2008). *Gabra2* encodes the alpha 2 subunit of the gamma-aminobutryic acid A (GABA-A) receptor, a pentameric, ligand-gated ion channel that mediates fast inhibitory neurotransmission via ligand-gated choloride influx and neuronal hyperpolarization (Olsen & Sieghart 2008). Gabra2-containing GABA-A receptors are expressed in several limbic regions involved in motivation and reward, including notably, the nucleus accumbens (Schwarzer *et al*. 2001; Morris *et al*. 2008). Chronic methamphetamine administration leads to an increase in Gabra2 mRNA in several brain regions, including the striatum and ventral tegmental area (Tamaki *et al*. 2008). Furthermore, *GABRA2* variants are associated with variance in psychostimulant behavioral responses and addiction, including cocaine (Dixon *et al*. 2010; Enoch *et al*. 2010; Smelson *et al*. 2012) and methyphenidate (Duka et al., 2015).

Interestingly, in contrast to our observation of enhanced methamphetamine behavior in B6J mice with the *Gabra2* intronic indel (corresponding to reduced Gabra2 expression), constitutive *Gabra2* knockout mice on a mixed C57BL/6J/129SvEv genetic background showed a decrease in cocaine-induced behaviors, including cocaine-induced reinforcement and locomotor sensitization (Dixon *et al*. 2010), with no phenotypic difference in cocaine intravenous self-administration or reinstatement of cocaine seeking (Dixon *et al*. 2014). Discrepancies could potentially be explained by multiple factors. First, the *Gabra2* intronic deletion is a different type of mutation compared to deletion of an entire exon (exon 4) in *Gabra2* knockouts (Dixon *et al*. 2008) – there is still a clearly detectable level of expression at the mRNA and protein level in multiple brain regions in B6J mice containing the *Gabra2* intronic indel (Mulligan *et al*. 2019). Thus, there could be fewer or different compensatory neuroadaptations in the GABA system (e.g., changes in other GABA-A subunits) in response to the constitutive *Gabra2* intronic indel compared to the *Gabra2* knockout. Interestingly, our findings are consistent with some of the literature. For example, like the *Gabra2* intronic indel, antisense oligodeoxynucleotides against *Gabra2* in the striatum adult Sprague-Dawley rats increased sensitivity to cocaine-induced locomotor activity and stereotypy (Peris *et al*. 1998). Second, methamphetamine has a different mechanism of action (reverses transport of monoamines) than cocaine (blocks transport of monoamines) and thus, it is possible that the *Gabra2* variants lead to different effects with different psychostimulants (Resnick *et al*. 1999). In support, the *Gabra2* locus was not identified to be linked to cocaine velocity in a reduced complexity cross that segregated this variant (Kumar *et al*. 2013). To our knowledge, behavioral responses to amphetamines have not been reported in *Gabra2* knockouts. Third, because the *Gabra2* knockout is on a mixed background containing 129SvEv and C57BL/6J alleles, including the *Gabra2* intronic indel, it is possible that phenotypic detection of exon 4 deletion is obscured by segregation of the *Gabra2* intronic indel and/or 129SvEv variants.

The impact of reduced Gabra2 levels on neuronal function has yet to be determined. Dixon and colleagues reported a 33% decrease in miniature inhibitory postsynaptic current (IPSC) and prolonged decay, but no difference in frequency in the nucleus accumbens of *Gabra2* knockout mice (Dixon *et al*. 2010). On the other hand, other groups have reported no change in CA1 pyramidal cell IPSCs following *Gabra2* genetic deletion (Panzanelli *et al*. 2011). Comparison of perisomatic asynchronous IPSCs in CA1 neurons between B6J containing the *Gabra2* intronic indel and the corrected B6J KI line at baseline and following treatment with a Gabra2/Gabra3 selective positive allosteric modulator (PAM) revealed a prolonged decay in the corrected KI line (Hawkins *et al*. 2021), indicating a restoration of PAM effects on receptor function. Kearney and colleagues found a reduction in *Gabra2*-containing receptors without a general reduction in overall GABA-A receptors in B6J versus corrected KI mice (Hawkins *et al*. 2021). The latter finding implicates a heteromeric receptor composition (containing both *Gabra1* and *Gabra2*) at perisomatic GABAergic synapses and that *Gabra2*-containing receptors might play a major role in mediating perisomatic phasic inhibition. These changes were associated with premature death and more severe seizures in B6J mice heterozygous for deletion of the Dravet syndrome candidate gene *Scn1a* relative to corrected lines heterozygous for the same deletion. However, inhibitory signaling was not profiled in other brain regions. Previous comparisons of GABA-A receptor mRNA expression in cortex, hippocampus, and striatum between B6J and the corrected KI line found a more profound alteration in subunit expression in the striatum compared to other brain regions (Mulligan *et al*. 2019). Taken together, these results suggest functional alterations in inhibitory signaling associated with a reduction of *Gabra2*, although the precise mechanisms and functional impact are likely to vary by cell type and brain region.

In the context of our previous findings (Mulligan *et al*. 2019), our current set of results support the notion that reduced *Gabra2* expression and plausibly, altered pentameric GABA-A receptor function leads to enhanced methamphetamine stimulant sensitivity which is in line with an inverse relationship between GABA-A receptor function (ability to transport chloride) and cocaine-induced locomotor stimulation (Peris 1996). To gain insight into neurobiological adaptations associated with reduced *Gabra2* expression, transcript covariance analysis of striatal tissue from F2 mice identified two other subunits that were positively correlated with Gabra2 expression (r ≥ +0.5, p < 0.05), including Gabre and Gabrg3 (**Table 3; Supplementary Table 4**) and two nominally correlated GABA-A subunit transcripts, including Gabrb1 (r = +0.37; p = 0.083) and Gabrq (r = +0.36; p = 0.089) (**Supplementary Table 4**). Thus, one hypothesis is that the concomitant decrease in expression in one or more of these GABA-A subunits with Gabra2 leads to decreased assembly and thus a decrease in the number of functional GABA-A receptors. In particular, Gabrg3 and Gabrb1 were highly expressed at a level comparable to Gabra2 (**Supplementary Table 4**), providing further confidence in these results. In support of Gabra2-Gabrb1 covariance in expression, a GABRA2 risk variant for alcohol dependence leading to reduced expression of GABRA2 positively correlated with expression of GABRB1, GABRG1, and GABRA4 in human iPSCs (Lieberman *et al*. 2015).

*Gabrb1* is coexpressed with *Gabra2* in vivo (Olsen & Sieghart 2008; Stephens *et al*. 2017). *GABRB1* variants have been associated with alcohol dependence and co-morbid substance use disorders (Stephens *et al*. 2017) and altered fMRI BOLD signal in multiple gyri and the caudate/insula during impulsivity and reward sensitivity tasks (Duka *et al*. 2017); thus, *GABRB1* is hypothesized to regulate excitability of GABA-A receptors in brain regions underlying reward-related behavior and possibly addiction. Concomitant decreases in Gabrb1 and Gabra2 expression were observed in cortex, striatum, and hippocampus of B6J mice relative to the corrected B6J KI line (Mulligan *et al*. 2019). In striatum, Gabra2 was previously found to assemble with Gabrb2 or Gabrb3 but not Gabrb1 (Hörtnagl *et al*. 2013), which could indicate additional changes in function of GABA-A receptors that do not contain Gabra2. Other subunits that were positively correlated with Gabra2 expression were expressed at a very low levels, including Gabre (r = +0.56; p = 0.0053) and Gabrq (r = +0.25; p = 0.089) (**Supplementary Table 3**). Thus, these correlations could be spurious.

Interestingly, in both F2 mice and *Gabra2* KI mice, the effect of *Gabra2* Genotype on acute methamphetamine-induced distance traveled was more pronounced in males versus females. The recapitulation of the sex-dependent genotypic effect from the QTL phenotype to the KI phenotype was striking (**Figs.3E,F vs. Fig.5B,C**) and further strengthens the support for the *Gabra2* indel as the causal, quantitative trait variant (Mulligan *et al*. 2019). What is the mechanism underlying the larger increase in methamphetamine-induced locomotor activity in males with the B6J *Gabra2* intronic deletion? One possibility is that male mutants show a larger reduction in Gabra2 transcript levels. Our present transcriptome dataset is not powered to detect sex differences; however, previous datasets fail to support sex-dependent effects of the B6J *Gabra2* indel on Gabra2 expression. We did not identify any previous sex differences in *Gabra2* expression between B6 substrains or in B6J relative to the *Gabra2*-corrected KI line (Mulligan *et al*. 2019). Furthermore, using a hypothalamic dataset from BXD strains on GeneNetwork that contained both sexes (Chesler *et al*. 2004; Mulligan *et al*. 2017), there was no sex difference in Gabra2 expression in BXD-RI strains with or without the B6J-derived Gabra2 intronic deletion as expression levels in female versus male BXD-Ri strains were highly correlated, regardless of *Gabra2* mutant genotype status (r = 0.91; p < 1 x 10^-16^; **Supplementary Figure 2**). Thus, sex differences in the mutational effect of the B6J-derived *Gabra2* intronic indel on transcription are unlikely to explain the enhanced effect of the chromosome 5 QTL or the *Gabra2* KI allele on methamphetamine sensitivity in males. Alternatively, sex differences in other neurotransmitter or neuromodulatory systems could interact with the *Gabra2* Genotype in determining the methamphetamine behavioral response. For example, sex-dependent interactions of endogenous opioid function and alcohol drinking on Gabra2 expression in mice have been reported (Rhinehart *et al*. 2018).

There are several limitations to this study. First, the parental B6 substrains and the F2 mice used for behavioral QTL analysis all had prior (but equal) exposure to naloxone over the course of 9 days that could have potentially influenced subsequent methamphetamine-induced behaviors as described in the Methods – this turned out not to be the case, as we nicely replicated the B6 substrain difference in methamphetamine-induced locomotor activity that was previously identified for methamphetamine-induced locomotor activity (B6J > B6N) (Kumar *et al*. 2013). Prior naloxone exposure could have enhanced our ability to detect the chromosome 5 QTL but this potential confound was offset by the fact that gene-edited *Gabra2* KI mice were completely naïve from any drug or experimental manipulations, yet the results qualitatively recapitulated the effect of the chromosome 5 QTL, despite the fact that the two studies were conducted in entirely different laboratories at different institutions. Another limitation is that the F2 mice from which samples were collected for striatal eQTL analysis had prior exposure to the mu opioid receptor agonist oxycodone as described in the Methods. However, this concern is mitigated by the fact that the eQTL was in the same direction as the prior report of Gabra2 expression differences between B6J and B6N (B6J < B6N) (Mulligan *et al*. 2019; Mortazavi *et al*. 2021). Finally, we only conducted eQTL analysis from a single brain tissue, the striatum. However, note that we previously identified reduced Gabra2 expression at both the transcript and protein levels in multiple brain regions, including cortex, hippocampus, and striatum (Mulligan *et al*. 2019); thus, the functional effects of this variant on Gabra2 expression are ubiquitous across CNS tissues examined.

To summarize, we replicated a historical QTL near the *Cyfip2* missense mutation underlying differential sensitivity to psychostimulant-induced maximum speed and identified a novel QTL and quantitative trait variant in *Gabra2* that underlies enhanced stimulant sensitivity to methamphetamine. This study further illustrates the efficiency of Reduced Complexity Crosses in systems genetic analysis of complex traits to rapidly identify causal genes and nucleotides (Bryant *et al*. 2018, 2020). Future studies will identify the brain regions, cell types, circuits, and physiological mechanisms underlying the relationship between reduced Gabra2 expression and GABA-A receptor function to enhancement of psychostimulant behaviors.

## DECLARATIONS

### CONSENT TO PUBLICATION

All authors have read and approved of the final version of the manuscript that has been submitted for publication.

### AUTHORS’ CONTRIBUTIONS

L.R.B., E.J.Y., and J.C.K. conducted a majority of the statistical analyses and data presentation and contributed to the writing of the manuscript. E.R.R and J.W.C. analyzed data. C.P., C.W., M.D., and S.K. generated data. J.A.B. and M.M.C. analyzed data. S.L.K. generated and analyzed data. G.E.H. generated reagents. R.W.W. contributed to writing the manuscript. C.D.B. conceived of the studies, advised in the analysis, and wrote the manuscript. M.K.M. generated and analyzed data and contributed to writing the manuscript.

## ACKNOWLEDGMENTS

We acknowledge the superb technical support of Sufiya Khanam and Christine Watkins (UTHSC).

## DATA AVAILABILITY STATEMENT

All data in its raw and processed forms will be made immediately available upon request.

---

**Supplementary Figure 1.**
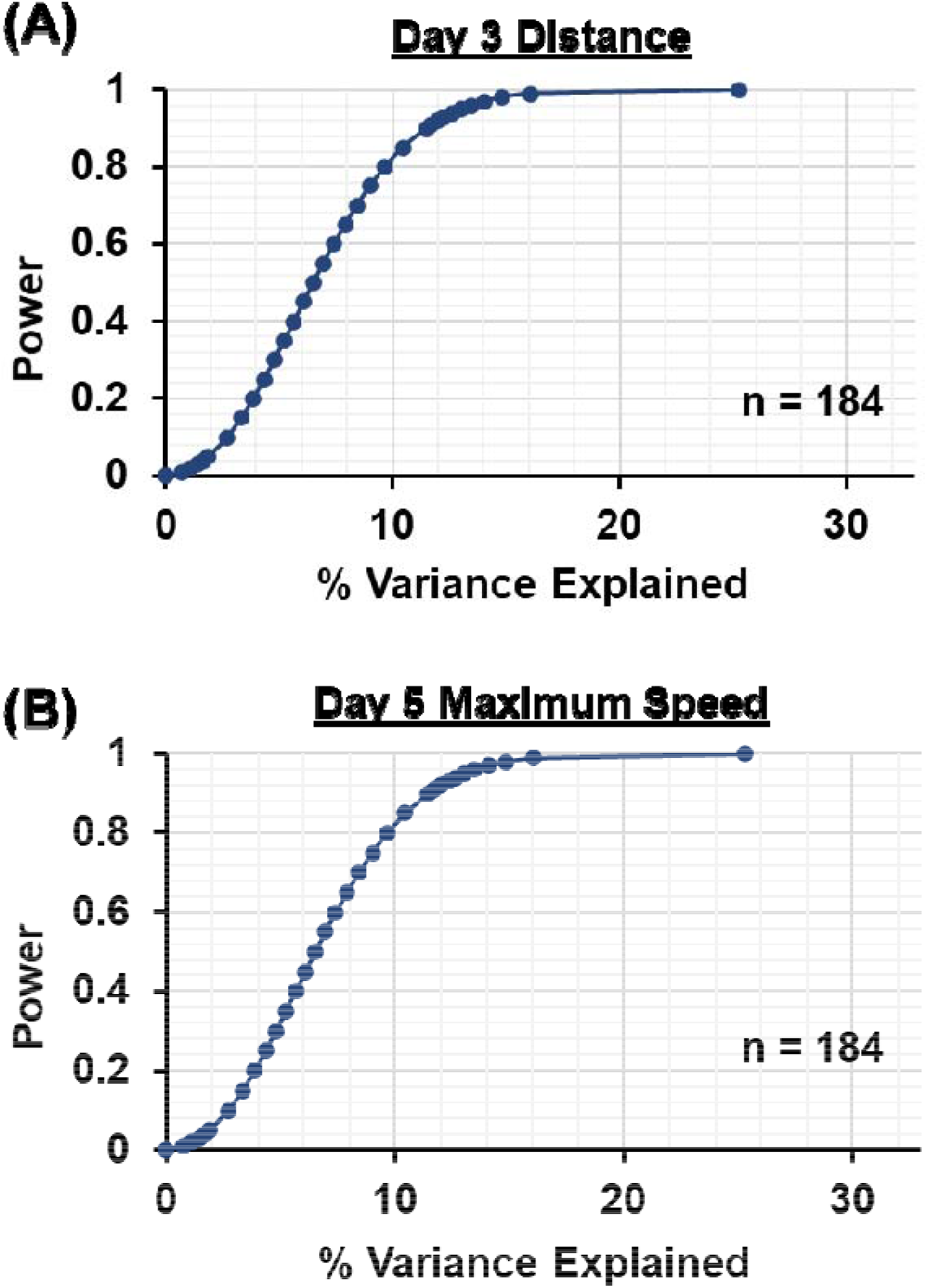
Power analysis for Day 3 distance traveled in B6J x B6NJ-F2 mice. Power versus effect size (% variance explained) for an additive QTL model (no covariates) and a sample size of 184 F2 mice. 0.2, 0.4, 0.6, and 0.8 power is achieved with an observed effect size of 3.86%, 5.65%, 7.41%, and 9.65% variance explained, respectively.

**Supplementary Figure 2.**
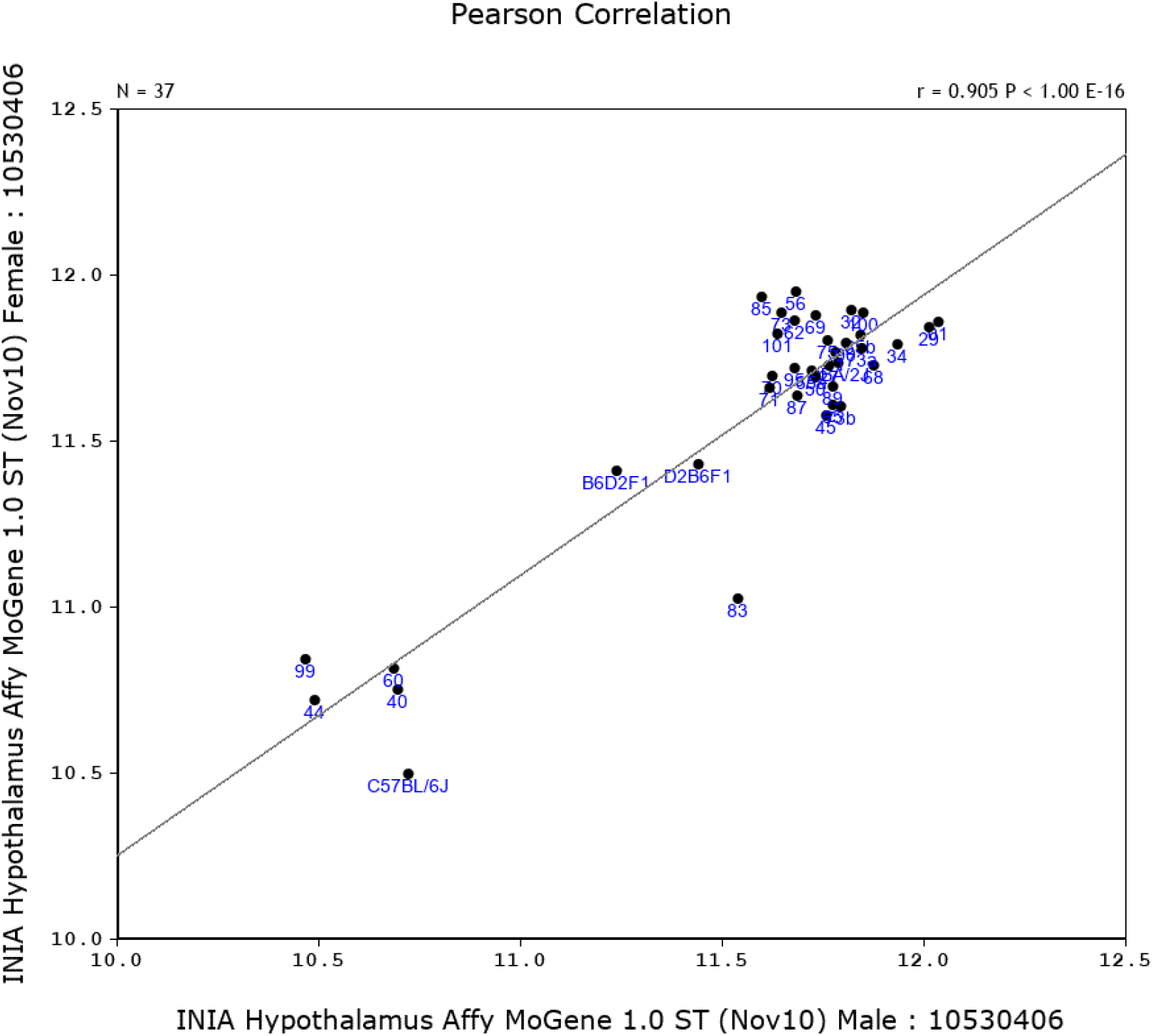
Correlation of Gabra2 expression in female versus male mice from various BXD-RI substrains with or without the *Gabra2* intronic indel. Plotted number indicate the BXD-RI strain. BXD-RI strains with the mutant Gabra2 allele include 40, 44, 60, 83, and 89. X and Y-axes indicate the dataset from GeneNetwork. The hypothalamus was selected for this analysis because it had both females and males contained within the dataset.

